# Potential Cancer Biomarkers: Mitotic Intra-S DNA Damage Checkpoint Genes

**DOI:** 10.1101/2024.09.19.613851

**Authors:** Kashvi Agarwal, Hengrui Liu

## Abstract

**BACKGROUND:** The mitotic intra-S DNA damage checkpoint signaling gene set is potentially involved in cancers where the genes play an important role. 17 total genes are involved in this gene set: ATF2, CHEK2, EME1, EME2, FANCD2, HUS1, HUS1B, MDC1, MRE11, MSH2, MUS81, NEK11, RAD17, RAD9A, RAD9B, TIPIN, XPC. The aim of this study is to complete a pan-cancer profile of each gene in the mitotic intra-S DNA damage checkpoint signaling gene set in order to determine potential diagnostic and prognostic purposes, while also determining how they could be used in a clinical setting as therapeutic targets to help patients.

**METHODS:** Multiomic data was acquired for the 17 genes; over 9000 samples of 33 types of cancer were analyzed to create pan-cancer profiles of CNV, mRNA expression, and pathway analysis.

**RESULTS:** The CNVs of some of these genes are associated with the survival of MESO, PCPG, BLCA, SKCM, LUAD, HNSC, LUSC, OV, and BRCA could be affected by the mRNA expression of the genes which can involve regulation of copy number.

**CONCLUSION:** With sufficient investigation, the genes involved in mitotic intra-S DNA damage checkpoint signaling may contribute to the development of cancer and may be used as biomarkers for cancer prognosis and diagnosis. To prove their clinical use for diagnosis and prognosis, however, and to create workable applications in clinical settings, further work is required. However, these pan-cancer profiles provide a more comprehensive knowledge of the mitotic intra-S DNA damage checkpoint signaling gene set in cancer as well as valuable information for future reference.

## Introduction

Cancer is defined as uncontrolled cell growth within the body, and it affects many people around the world. It is such a major disease that approximately 2 million people are expected to be diagnosed with it in 2024 and about 611,720 will die from it. When looking at specific cancers, the most common cancer diagnosis is breast cancer and second is prostate cancer. On the other hand, most cancer deaths result from lung and bronchus cancer with 125,070 people expected to die in 2024 from them [1]. Cancer occurs as the cell division is faulty and cells continue dividing. This can result in tumors which can be benign or malignant. Benign tumors are typically those that do not spread or invade nearby tissue and can sometimes be removed without the risk of recurrence. Malignant tumors (cancerous tumors) spread and invade nearby tissue through metastasis and can be solid of liquid. Cancers are typically a genetic disease which can occur due to errors during cell division, damaged DNA, or inheritance. The human body has the ability to kill cells with damaged DNA, but this process is slowed down as age increases, making cancer more prevalent in later life [2, 3].

Completing a pan-cancer analysis allows for comparisons within a specific gene set across most cancer types and create similarities and differences. This type of analysis helps with having early diagnoses and prognoses which could potentially save more lives.

An important aspect of cancer development is unregulated mitosis (cell division). Under perfect conditions, mitosis is supposed to yield two cells with the correct number of chromosomes. However, when mitosis malfunctions, a cell could end up with an abnormal chromosome number (aneuploidy), which could potentially be cancer [3, 4].

Genetic testing can greatly help people who have a history of cancer in their family, or they inherited a gene mutation. Once a person has been diagnosed with cancer, certain tests on cancer cells can help locate protein or gene changes. It becomes a major tool when it can help determine a person’s future prognosis and which treatments are best suitable [5]. In this way, testing in cancer related gene sets puts the field at an advantage of early diagnosis with better outcomes.

The gene set analyzed from the Gene Ontology (GO) database[6] is called the mitotic intra-S DNA damage checkpoint signaling and has the following genes involved in its pathway: ATF2, CHEK2, EME1, EME2, FANCD2, HUS1, HUS1B, MDC1, MRE11, MSH2, MUS81, NEK11, RAD17, RAD9A, RAD9B, TIPIN, XPC. This set is a mitotic cell checkpoint that slows DNA synthesis in response to DNA damage by the prevention of new origin firing and the stabilization of slow replication fork progression [5]. When disrupted, this checkpoint will stop functioning and allow for a continuation of DNA synthesis disregarding the replication of damaged DNA, which is a large part of cancer growth within the body[7, 8].

Within this gene set, certain genes such as ATF2 and CHEK2 are positive regulators in terms of DNA repair[9] [10, 11]; others such as FANCD2 and MRE11 are negative regulators in terms of DNA repair[12, 13]. Understanding specific roles of each gene plays a crucial role in understanding this gene set as a whole and advancing the field of science by diagnosing cancer earlier with this gene set as a new biomarker.

Generally, biomarkers serve a large purpose in the field of research and diagnosis as they facilitate a deeper comprehension of disease processes and the mechanisms by which medications function to treat illness[14]. With this understanding, illness can be prevented before it manifests or diagnosed early. Biomarkers can be utilized in the development of novel medications as well as to increase the safety and effectiveness of current ones. Especially for cancer, biomarkers help doctors treat certain types of cancers and provide the correct treatment without inflicting more harm or pain.

We analyzed these 17 genes across over 9,000 samples from 33 distinct cancer types using multi-omic profiling data from TCGA, a resource widely utilized in numerous previous studies[15-33]. This extensive pan-cancer analysis provided an overview of these genes for future reference by revealing the many expression controls of the genes within the mitotic intra-S DNA damage checkpoint signaling gene collection and examining possible correlations with widely used cancer signaling pathways. These profiles, in our opinion, will yield genetic data that will be beneficial for further research on cancer.

## Methods

### Data Acquisitions

All data for the RSEM normalized mRNA expression data, copy number variant (CNV), and pathway activity were obtained from The Cancer Genome Atlas Database (TCGA).

### Copy Number Variation Analysis

The CNV summary pie chart take data of 11495 samples and processed through GISTIC2.0. This allowed identification of important altered regions of deletion or amplification across all sets of patients.

To understand the relationship between CNV and mRNA expression, mRNA expression and CNV data were normalized by RSEM and then downloaded from the TCGA database. The TCGA barcode then merged the mRNA expression data and CNV raw data. The Spearman correlation analysis was done to obtain the correlation between CNV and mRNA expression. The p-value was then adjusted by FDR. For CNV and survival 11495 samples were downloaded and uncensored data was left out. Those with a competing risk of death because of cancer were removed.

Clinical survival data and CNV data were merged by sample barcode. These samples were then split into Amp., Dele., and WT groups. R package survival was used to fit survival time and status within the two groups and Logrank tests were done to understand survival difference between groups.

### Expression Analysis

The differential expression was based on corrected RSEM mRNA expression. The fold change was calculated by the mean of Tumor / mean of Normal and the p-value was calculated by the t-test and finally adjusted by the FDR.

To identify variations in gene expression that are important to a subtype, the expression and subtype analysis was utilized. Clinical data from tumor samples from nine different cancer types (HNSC, LUSC, COAD, STAD, LUAD, GBM, BRCA, KIRC, and BLCA) were employed by GSCA for this function. By using the sample barcode, data on clinical subtype and mRNA expression were combined. There must be a minimum of 5 samples in the subtype subgroup. Using the Wilcoxon test (number of subtype groups == 2) and the ANOVA test (number of subtype groups > 2), GSCA compared the GSVA score between groups.

For expression and survival, clinical data from tumor samples were used and those with a competing risk of death because of cancer were removed. A median mRNA value allows tumor samples to be split into low and high expression group, and then R package survival was used to fit survival time and status within the two groups. Logrank tests and Cox Proportional-Hazards model were done for every gene in every cancer.

### Pathway Analysis

This analysis looks at and estimates the difference between pathway activation and inhibition in gene expression as defined by median pathway scores.

For 7876 samples (from the TCGA database) representing 32 different cancer types, the pathway activity score of 10 cancer-related pathways was determined using reverse phase protein array (RPPA) data from the TCPA database. Western blot processes are comparable to those of RPPA, a high-throughput antibody-based approach. After being removed from tumor tissue or cultivated cells, proteins are denatured by SDS and printed onto slides covered with nitrocellulose, which is followed by an antibody probe.

TSC/mTOR, RTK, RAS/MAPK, PI3K/AKT, Hormone ER, Hormone AR, EMT, DNA Damage Response, Cell Cycle, and Apoptosis pathways are among the pathways that the GSCA covered. These are all well-known pathways linked to cancer.

To determine the relative protein level for each component, RBN RPPA data were median-centered and standardized by the standard deviation for all samples. The total of all positive regulatory components’ relative protein levels less all negative regulatory components’ relative protein levels in a given route is the pathway score.

The median gene expression of the samples was used to split them into two groups (High and Low). The student T test was used to determine the difference in pathway activity score (PAS) between the groups, and FDR was used to modify the P value. A FDR of less than 0.05 is regarded as significant. Gene A may have an activating influence on a route when PAS (Gene A High expression) > PAS (Gene A Low expression), and it may have an inhibiting effect on a pathway otherwise.

## Results

### Copy Number Variation Analysis

Copy number variation (CNV) profiles were analyzed for this gene set. This analysis showed a variety of patterns crucial to understanding the gene set across multiple cancer types.

The CNV pie chart and plots displayed the large abundance of heterozygous and homozygous CNV, however, there seems to be more heterozygous CNV than homozygous CNV when look at the pan-cancer analysis (Figure 1).

**Figure 1:**
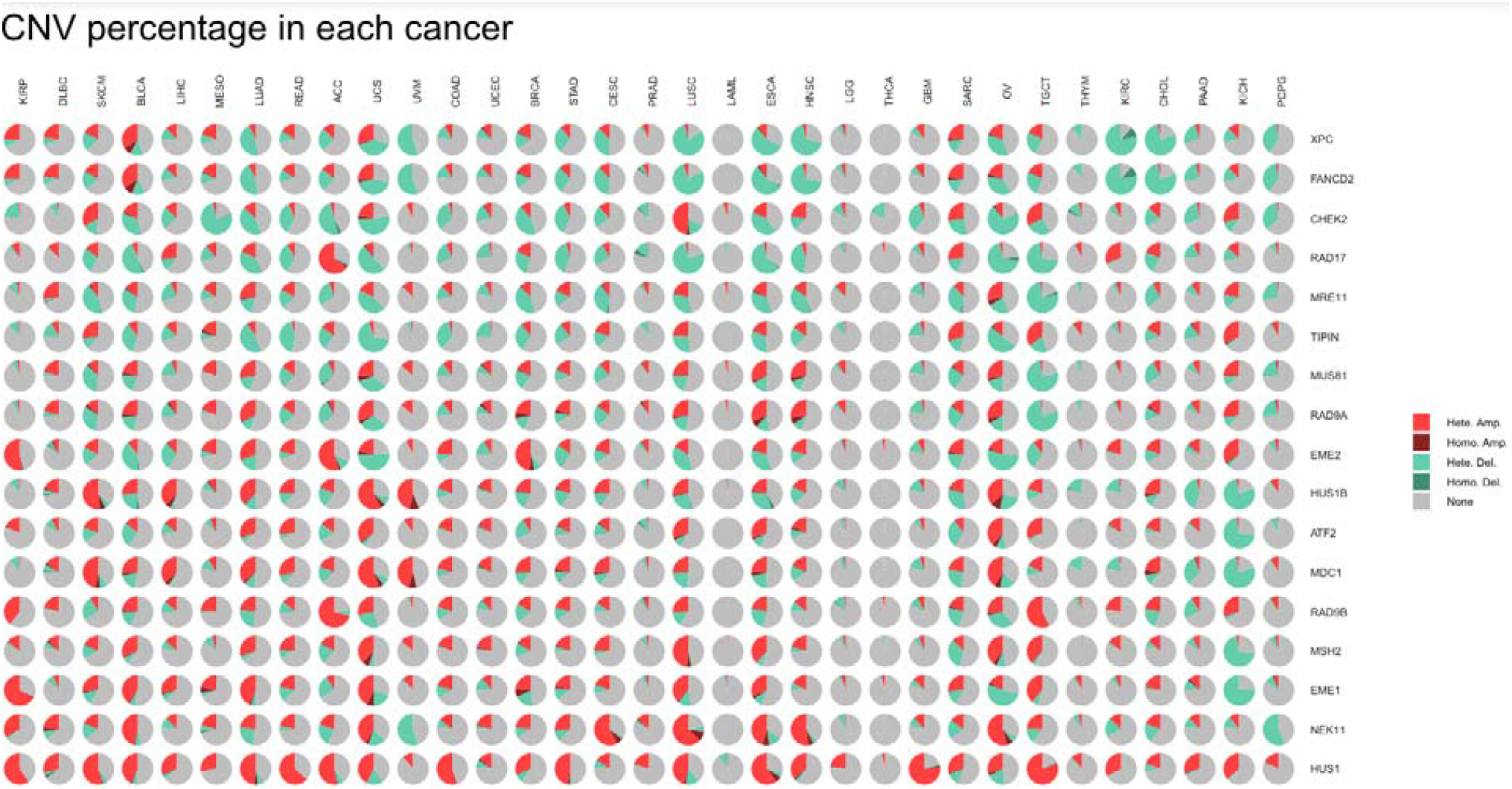
Pie charts of CNV distribution across cancers. Hete Amp = heterozygous amplification; Homo Amp = homozygous amplification; Hete Del = heterozygous deletion; Homo Del = homozygous deletion

The CNV pan-cancer survival analysis portrayed the importance that many genes within this gene set are associated with patient survival. The specific cancer types that these genes are most associated with include KIRP, MESO, PCPG, LIHC, and PRAD. The genes associated with cancers MESO, PCPG, and PRAD for the overall survival include MUS81, XPC, and HUS1, respectively. In terms of the disease specific survival, gene NEK11 is associated with KIRP, genes MUS81 and RAD9A are associated with MESO, XPC is associated with PCPG, gene MDC1 is associates with LIHC, and HUS1 is associated with PRAD (Figure 2)

**Figure 2:**
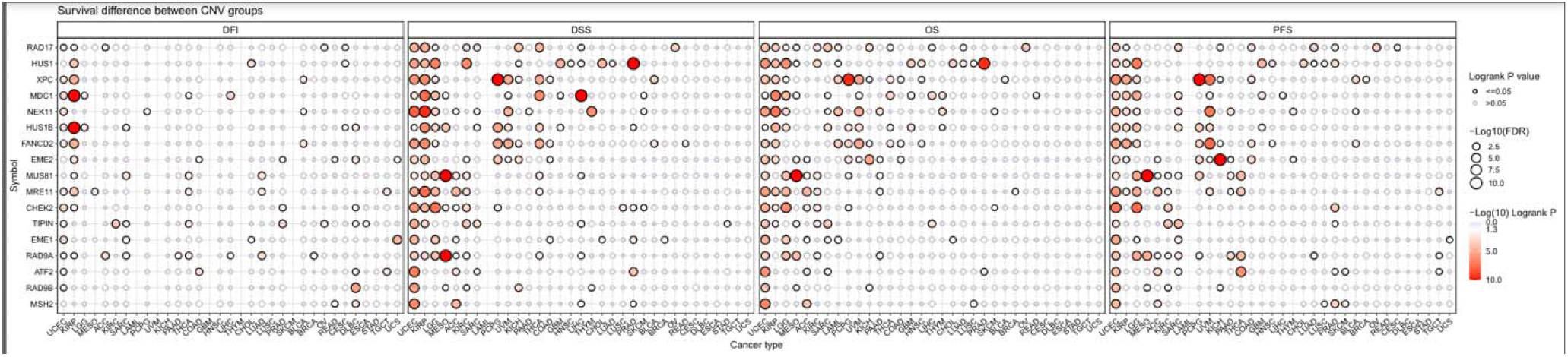
Survival difference between CNV groups.

When looking at the correlation between CNV and mRNA expression, it is seen that CNV is positively correlated with most of the genes in the set by the Spearman Correlation.

Because of the results from the Copy Number Variation profiles, it can be said that there is a potential association of the heterozygous CNV of the genes within the mitotic intra-S DNA damage checkpoint signaling gene set and cancers (Figure 3).

**Figure 3:**
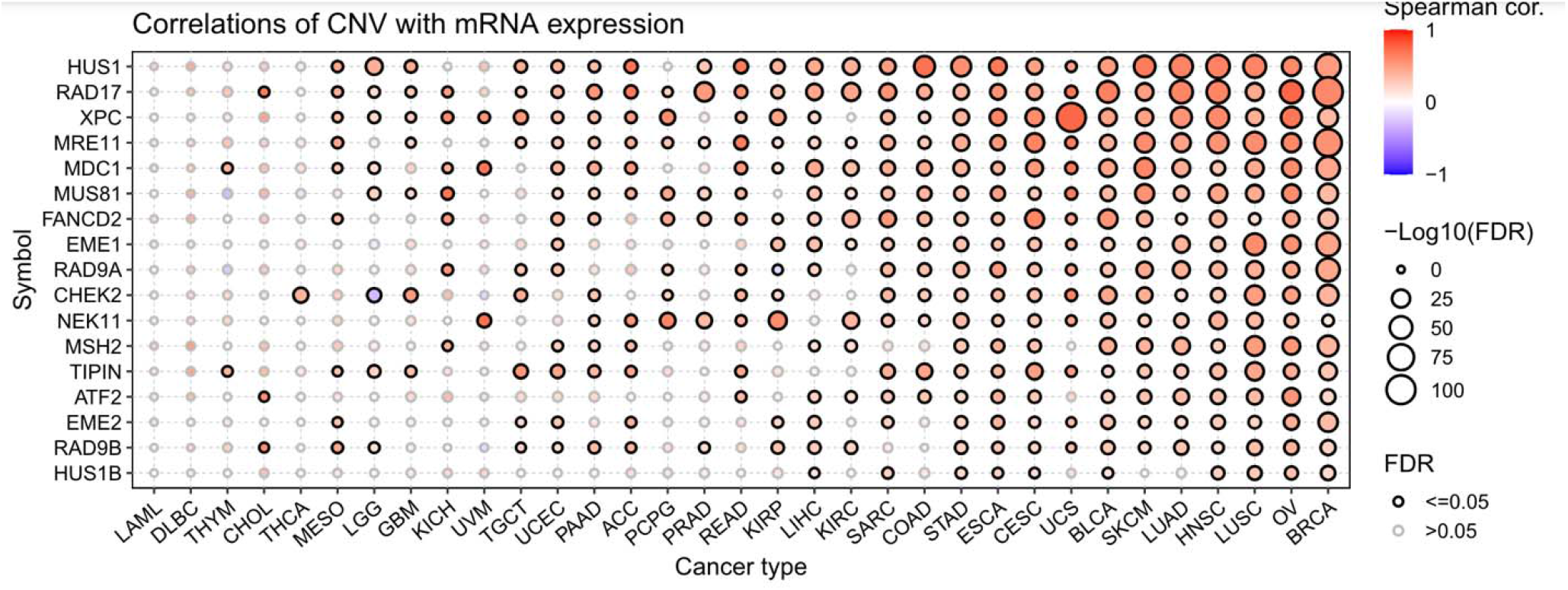
Correlation plot between CNV and mRNA expression with the spearman correlation as the correlation coefficient.

### mRNA Expression Analysis

The expression profiles of the gene set were examined pan-cancer. Data was used to analyze a difference in mRNA expression between cancer and non-cancer tissues of the genes within the mitotic intra-s DNA damage checkpoint signaling. Based on the data, the genes EME1, FANCD2, CHEK2, MSH2, and TIPIN all seem to be upregulated in the cancer LUSC.

The most significant appear to be EME1 and CHEK2, however EME1 and FANCD2, appear to be the most significant in other cancers as well, which include LUAD, BRCA, LIHC, and LUSC.

Moreover, the genes NEK11 and XPC appear to be downregulated in cancers KICH and LUSC, respectively. Out of the two, gene NEK11 had a higher fold change. From a broader perspective, genes EME1 to HUS1, going down the gene symbol key, were upregulated in most cancer, while genes MDC1 to NEK11 were downregulated in some cancers (Figure 4).

**Figure 4:**
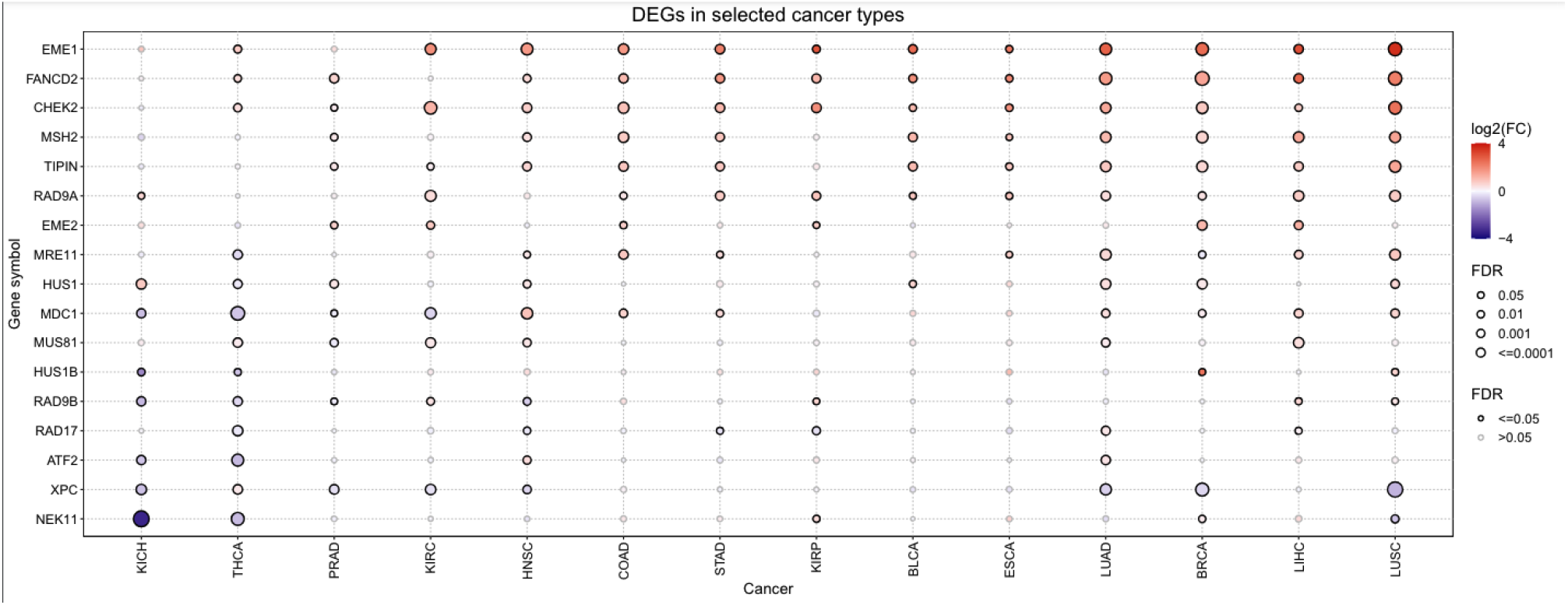
Expression difference between non-cancer and cancer tissues.

The subtype difference between high and low gene was also analyzed. Cancers BRCA and KIRC displayed to be the most significant between the genes within the set. STAD was also seen to have subtype significance, but less than the other two when looking at the genes set as a whole (Figure 5).

**Figure 5:**
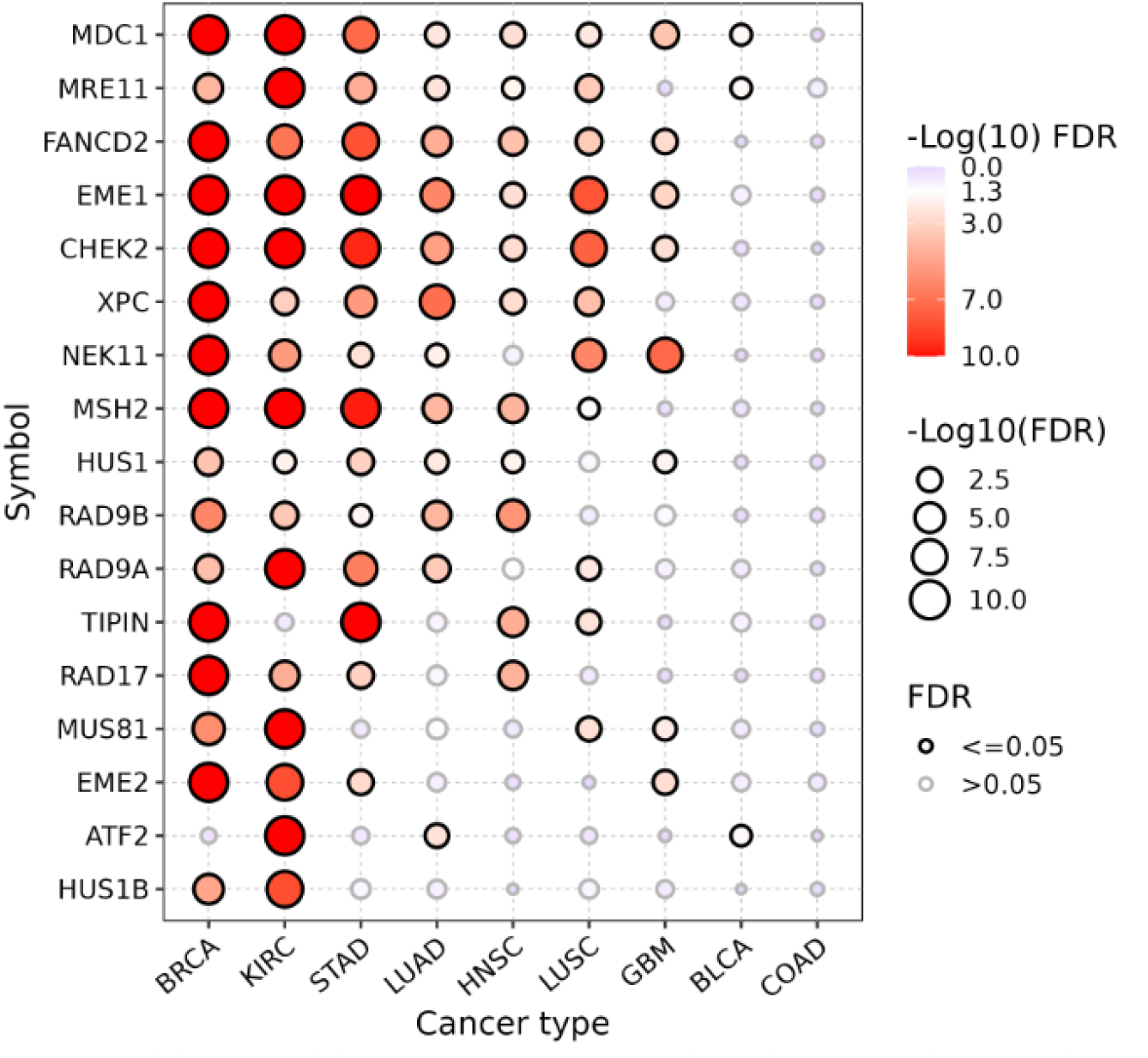
Difference of expression between subtypes of cancers.

By looking at the survival difference between high and low gene expression, it was established that certain genes show a relationship between expression and survival. Genes EME1 and CHEK2 show a larger higher survival risk when overexpressed in cancer ACC (Figure 6).

**Figure 6:**
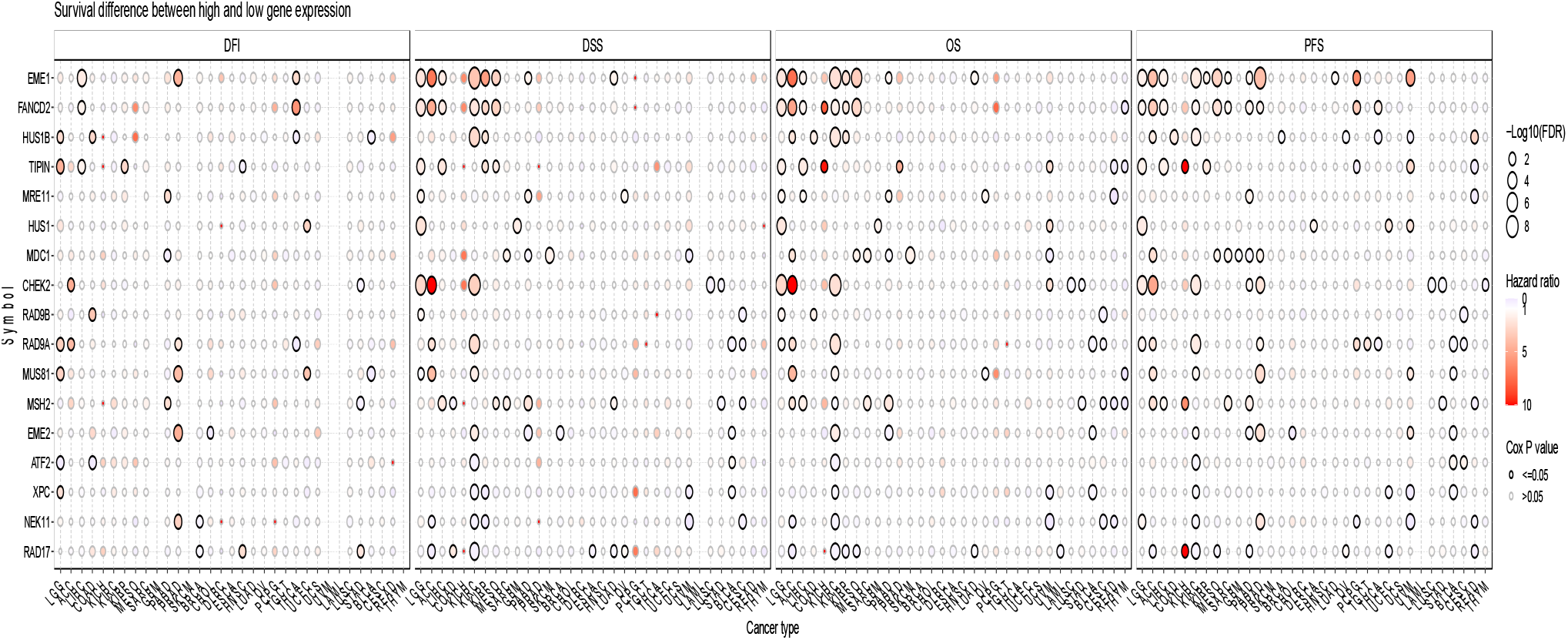
Survival difference between high and low gene expression.

### Pathway Activity Analysis

Higher expression of EME1 in 72% of cancer types analyzed was associated with cell cycle activation. FANCD2 had a similar association within 66% of the cancer types analyzed, as well as did TIPIN for 53% of cancer types. Also, XPC and NEK11 are associated with both apoptosis inhibition and cell cycle inhibition. MDC1, FANCD2, and EME1 are seen to have a relationship with the inhibition of the hormone estrogen receptor. Genes TIPIN, MUS81, MSH2, FANCD2, EME1, and CHEK2 relate to the inhibition of RASMAPK between cancer types (Figure 7).

**Figure 7:**
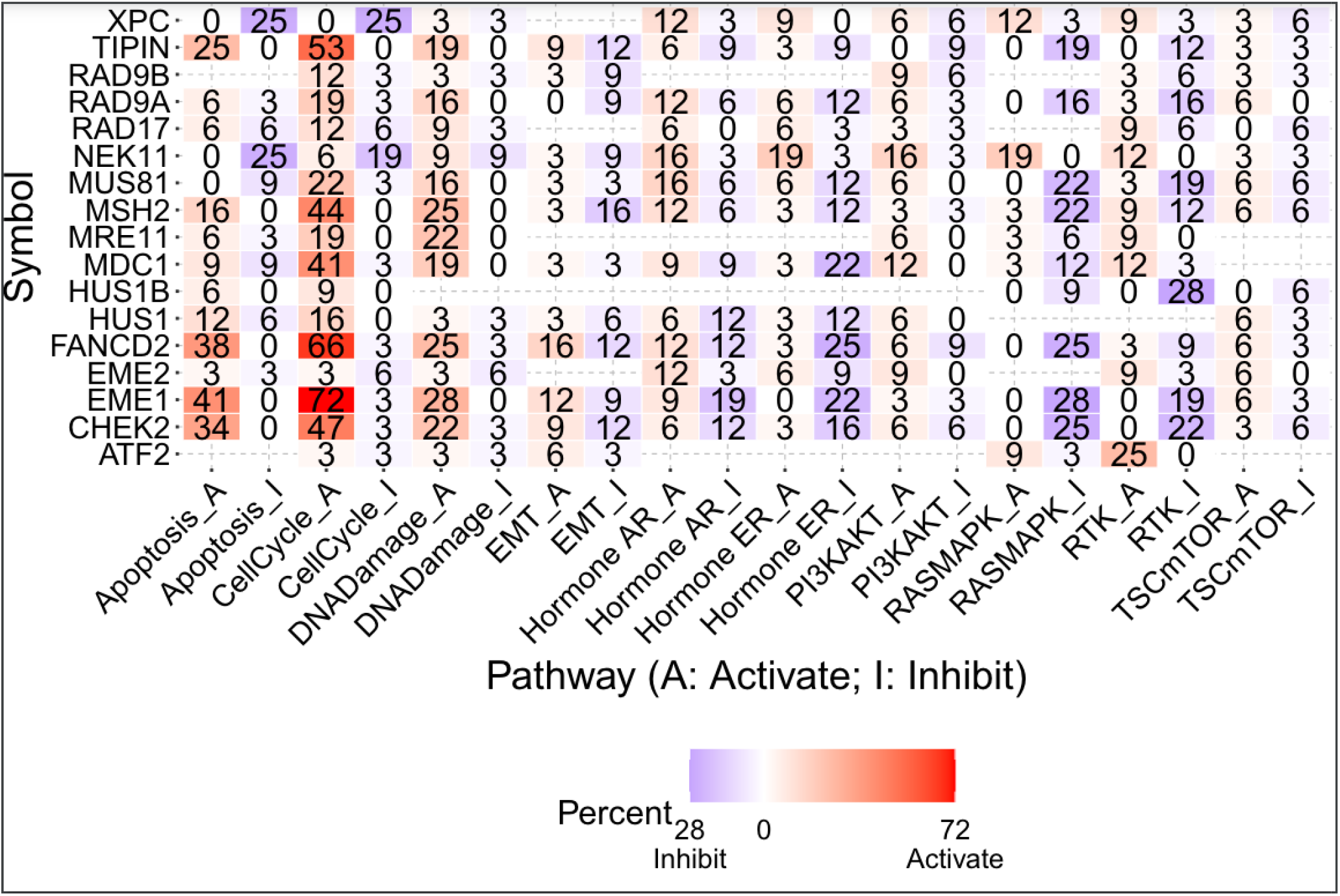
Pathway activity and expression of the genes within gene set.

## Discussion

Cancer research is a highly active field that has been extensively explored in numerous studies[34-43]. The pan-cancer analysis of the genes within the mitotic intra-S DNA damage checkpoint signaling set has portrayed themselves to be valuable in terms of identifying cancer diagnostic and prognostic biomarkers. Within each analysis, we performed various correlations in order to account for multiple effects of the gene on various systems part of our bodies.

One of our Copy Number Variation Analysis’s, specifically overall survival, displays that gene MUS81 heavily impacted the survival time of MESO patients. It was able to increase their time as it is a known tumor suppressor. Previous studies [44] have found results accordingly to ours, as they state that a deficiency or insufficiency of MUS81 caused a spike in cancer susceptibility. This study tested on mice specifically and found that lower levels of MUS81 lead to sarcoma development, showing the importance that this gene plays on tumor suppression. This gene could be a potential biomarker for cancer type MESO, however, our results from the other analysis do not provide enough information for the assertion, so future research on this gene is recommended.

Within the CNV analysis, we also looked at the correlation between CNV and mRNA expression. The results showed that most of the genes from the set showed a positive correlation with cancer types BLCA, SKCM, LUAD, HNSC, LUSC, OV, and BRCA, which the most overall positive correlation being with cancer BRCA. In this case, the genes that were the most positively correlated with cancer type BRCA were RAD17 and MRE11. Previous studies and databases [45] have classified gene RAD17 as being involved in breast tumorigenesis and its turnover is important for continued genome stability and carcinogenesis. Also, MRE11 has been defined by various research [13] as being associated with several cancer types, including breast cancer, when its amount is varied. Higher MRE11 levels have shown to be link to lower survival rates in patients receiving treatment for breast cancer. In this case, both RAD17 and MRE11 have portrayed themselves to be biomarkers for a few cancer types, especially BRCA, and this statement is also supported by various research on these genes.

Our study also focused on a mRNA expression. Analysis, first looking at the expression difference between non-cancer and cancer tissues. This analysis allowed us to understand the importance certain genes play when downregulated or upregulated on cancer. Specifically, gene NEK11 was seen to be greatly downregulated in cancer KIHC, meaning it decreased production of cellular components (in our case, of tumors) [46]. Various studies have also defined this gene as a candidate for strongly reducing proteins caused by degradation, proving its overall effect on cancers like KIHC[47]. This being said, downregulation of gene NEK11 is a potential biomarker for possible tumor growth as it has been working to decrease the mass of the tumors, but further research is still recommended. Furthermore, our results depicted genes that were heavily upregulated in cancer type LUSC, which were EME1 and CHEK2. Upregulation of these genes means that they were essentially increasing tumors and can be defined associated with cancer initiation and progression [48]. For gene EME1, previous studies have agreed with our results, describing it as being upregulated and strongly associated with lymph node metastasis [49]. This shows that EME1 can be a potential biomarker for cancer type LUSC, making it valuable to research. Gene CHEK2 was also upregulated, but previous research suggests that CHEK2 is more dangerous when it has been mutated, and the mutation can lead to an increased risk of cancer [10, 50]. Because our study did not look at the effects of mutated genes on cancer, an explanation of gene CHEK2 cannot be provided from us and more research is suggested.

Our next mRNA expression analysis looked at the difference of expression between subtypes of cancers. Cancers BRCA and KIRC displayed to be the most significant between the genes within the set. STAD was also seen to have subtype significance, but less than the other two when looking at the genes set as a whole. Our results are not able to say much more regarding this analysis, so further research is needed to accurately determine how the genes in our gene set can be potential biomarkers for a diagnosis/prognosis.

The final mRNA expression analysis we conducted was the survival difference between high and low gene expression. Genes EME1 and CHEK2 show a larger higher survival risk when overexpressed in cancer ACC. These genes have already been defined as biomarkers for diagnosis and prognosis in cancer LUSC and by previous research, so they can be further defined as biomarkers for cancer ACC. These 2 genes played an important role as they were upregulated and increased tumor mass within the body. As a result, they show an increased survival risk among patients because those patients have major tumor growth within their bodies.

The pathway analysis was the last analysis we used, and we specifically looked at the pathway activity and expression of the genes within gene set. The most notable gene was EME1, as it was observed to active the cell cycle pathways in 72% of the samples cancer types. Previous research also states that EME1 is notable, as it is highly upregulated and causes progression of many cancer types [50]. Because it could be a potential biomarker, further research is highly recommended to understand this connection of cell cycle activation by EME1 and its upregulation.

Taking all of this into account, our study does have a few limitations that could affect the results. First, only tissue samples were analyzed, meaning they we were not able to understand the cancers as they evolve and change. Many environmental factors also affect gene relationships (drug treatments, presence of mutagens, microbiotics, etc.), proving to be a limitation[51]. Our research was also limited by the extent of analysis, making it worthwhile to conduct future research on this gene set to establish a definitive conclusion. Our study does provide valuable information by providing possible diagnostic and prognostic biomarkers. For future use as a guide in cancer research, these profiles also provide a general overview of the genes in the mitotic intra-S DNA damage checkpoint signaling gene set.

## Conclusion

This study provided valuable insight into CNV, mRNA expression, and pathways crosstalk profiles for the genes associated with the mitotic intra-S DNA damage checkpoint signaling gene set across 33 different cancer types. The profiles might help comprehend future cancer treatments as possible prognostic and diagnostic biomarkers,, such as EME1 in ACC and LUSC. With sufficient investigation, the genes in the mitotic intra-S DNA damage checkpoint signaling gene set may contribute to the genesis of cancer and may one day be used as biomarkers for cancer prognosis and diagnosis. To prove their clinical use for diagnosis and prognosis, however, and to create workable applications in clinical settings, further work is required. However, these pan-cancer profiles provide a more comprehensive knowledge of mitotic intra-S DNA damage checkpoint signaling in cancer as well as valuable information for future reference.

## List of the cancer-type abbreviations

ACC: Adrenocortical carcinoma
BLCA: Bladder Urothelial Carcinoma
BRCA: Breast invasive carcinoma
CESC: Cervical squamous cell carcinoma and endocervical adenocarcinoma
CHOL: Cholangio carcinoma
COAD: Colon adenocarcinoma
DLBC: Lymphoid Neoplasm Diffuse Large B-cell Lymphoma
ESCA: Esophageal carcinoma
GBM: Glioblastoma multiforme
HNSC: Head and Neck squamous cell carcinoma
KICH: Kidney Chromophobe
KIRC: Kidney renal clear cell carcinoma
KIRP: Kidney renal papillary cell carcinoma
LAML: Acute Myeloid Leukemia
LGG: Brain Lower Grade Glioma
LIHC: Liver hepatocellular carcinoma
LUAD: Lung adenocarcinoma
LUSC: Lung squamous cell carcinoma
MESO: Mesothelioma
OV: Ovarian serous cystadenocarcinoma
PAAD: Pancreatic adenocarcinoma
PCPG: Pheochromocytoma and Paraganglioma
PRAD: Prostate adenocarcinoma
READ: Rectum adenocarcinoma
SARC: Sarcoma
SKCM: Skin Cutaneous Melanoma
STAD: Stomach adenocarcinoma
TGCT: Testicular Germ Cell Tumors
THCA: Thyroid carcinoma
THYM: Thymoma
UCEC: Uterine Corpus Endometrial Carcinoma
UCS: Uterine Carcinosarcoma
UVM: Uveal Melanoma

## Declarations

## Author Contributions

Kashvi Agarwal conducted all analyses and drafted the manuscript. Hengrui Liu designed the study, supervised the project, provided guidance to Kashvi Agarwal, and edited the manuscript.

## Acknowledgments

This study was conducted during the Lumiere project and received support from the Lumiere Company.

## Availability of data and materials

The source of the raw data was provided in the paper and the raw analysis data of this study are provided by the corresponding author with a reasonable request.

## Ethical approval

Not applicable.

